# Novel role of *Periploca laevigata* extracts as anti-diabetic and anti-inflammatory function in pancreatic β cells exposed to hyperglycaemia

**DOI:** 10.1101/2023.11.07.565784

**Authors:** Ghada Trabelsi, Susana Mellado, Zahar Kalboussi, Leila Chekir-Ghedira, María Pascual

## Abstract

Type 2 Diabetes (T2D) is a major health problem worldwide. This metabolic disease is associated with high blood glucose levels due to insufficient insulin production by the pancreas, leading to an inflammatory immune response. Considering the beneficial roles of medicinal plants to control the diabetic complications of T2D, as well as the therapeutic roles of the extracts from *Periploca laevigata* (PL), this study evaluates the anti-diabetic and anti-inflammatory properties of PL extracts in the pancreatic β cell line INS-1E under hyperglycaemic. Our findings demonstrated, for the first time, that whereas PL extracts tends to upregulate the insulin gene expression, PL extract from leaves increase significantly the gene expression of GLUT-2 and the transcription factor PDX-1 in glucose-treated INS-1E cells. Notably, some PL extracts are also capable to decrease significantly the gene expression of iNOS and to increase the IL-10 gene expression in glucose-treated INS-1E cells. However, the PL extracts show a tendency to decrease the gene expression of NF-κB and MCP-1 in glucose-treated INS-1E cells. In conclusion, our findings suggest that the treatment with PL extracts could benefit the pancreatic β cell function and alleviate the pancreatic inflammation, thereby representing potential advantages of the glucose metabolism in the T2D.

## INTRODUCTION

Among the unresolved pathologies until today that cause severe health problems and which is steadily increasing is type 2 diabetes (T2D). Worldwide, the incidence of diabetes mellitus is increasing at an alarming rate, and it is predicted to accelerate further and reach epidemic proportions [1]. One of the main causes of the T2D is reduced insulin secretion due to pancreatic β-cell dysfunction [2]. Pancreatic beta cells secrete insulin in response to glucose stimulation to maintain blood glucose levels within a relatively narrow range [3]. Insulin is required to transport glucose from the bloodstream to target tissues. High blood glucose levels in the body are able to cause problems associated with insulin secretion, insulin action, or both [4]. In case of people suffer this disease, insulin can be prescribed as a first-line treatment for those intolerant to other antidiabetics drugs or in individuals with a primary β-cell defect [5].

T2D is a dual-etiological, inflammatory and metabolic disease [6]. This inflammatory process contributes to loss of metabolic tolerance and increased risk of insulin resistance. Indeed, recent reports showed that the use of antagonists against inflammatory molecules (e.g., IL-1β, IL-6, TNF) is capable to improve insulin secretion and blood sugar regulation [7–9]. Thus, therapeutic interventions counteract metabolic inflammation, improve insulin secretion and action, control blood sugar, and can prevent long-term complications [10]. However, diabetes is poorly treated. While the most commonly used drugs (e.g., metformin and sulfonyles) do not cause lasting control of diabetes, insulin and other therapies (e.g., thiazolidineddiones, inhibitors of glucosidases) have adverse effects [11–13]. In this context, several countries use medicinal plants, which play a role in the management of T2D, by delaying the development of diabetic complications and the correction of metabolic abnormalities [14].

*Periploca laevigata* (PL) belonging to family *Apocynaceae* [15], is native to Mediterranean region and widely distributed in the Sahara area [16]. This shrub grows mainly in semi-arid and sub-humid areas, and contains various pharmacological activities such as antioxidant and antimicrobial effects [16,17]. Previous studies demonstrated that the pretreatment with the PL extract from seeds has an hepatoprotective, anti-inflammatory and analgesic properties [18]. Studies on the chemical constituents have shown the presence of flavonoids, polyphenols, proteins, triterpenic compounds and oleanolic acid derivatives [18,19]. For instance, oleanolic acid and triterpenic compounds have important pharmacological functions, such as anti-bacterial activities [19]. The novel polysaccharide named PLP1 from root barks exhibits antioxidant and antibacterial activities [20].

Considering the beneficial roles of medicinal plants to control the diabetic complications of T2D, as well as the therapeutic role of PL extract, this study evaluates the anti-diabetic and anti-inflammatory properties of PL extracts in a widely used pancreatic β cell line. Our findings demonstrated for the first time the potentially beneficial effects of the PL extracts on the expression of insulin- and inflammatory-related genes in INS-1E cells under high glucose conditions.

## MATERIALS AND METHODS

### Plant material

Root, fruit and leaves parts of *Periploca laevigata Ait* (also named *Periploca angustifolia Labill*) was collected from Soukrine (Monastir,Tunisia, LAT: 35.6528/LON: 10.9592) on June 2021. Identification was carried out by Pr. Skhiri Fathia (Higher Institute of Biotechnology of Monastir, Tunisia), according to the flora of Tunisia. The registered specimen has been kept in the Laboratory of Pharmacognosy, Faculty of Pharmacy of Monastir, Tunisia. The raw material (root, fruit and leaves) was washed with distilled water and then dried at room temperature for at least 30 days. The dried root, fruit and leaves were further crushed to obtain a fine powder, and then stored in glass bottles at room temperature.

### Isolation and purification of plant extracts

*Periploca laevigata* root (PLR), fruit (PLF), and leaves (PLL) powder (100 g) were boiled with distilled water (1000 mL) for 20 min, and then the filtrate was drying by lyophilization, and stored at -4□°C for further use.

In addition, we also isolated PLL by using a hydro-ethanolic solution (PLL-E). PLL powder (100 g) was extracted with 1 L of 70% ethanol. The filtrate was concentrated by a rotary evaporator under reduced pressure approximately at 78.4□°C. Then, it was dried by lyophilization to produce the crude extract, and stored at -4□°C for further use.

### SPME-GC-MS analyses

Volatile fruit aroma constituents from PL were analyzed by using the solid-phase microextraction (SPME) coupled to gas chromatography mass spectrometry (SPME-GC-MS). It allows the extraction and concentration of compounds which are found in few quantities of liquid. The samples were conditioned in thermostatic bath at 70.0 ± 1.0 ºC for 1 h. Then, sample/headspace equilibration period, the septum covering the vial headspace was pierced with the needle containing the fiber retracted for 30 min at 22.0 ± 1.0 ºC. Supelco (Bellefonte, PA) SPME Fibre 100 μm Polydmethylsiloxane (PDMS) was used to sample the free space of the extracts inserted into a 5 mL septum glass. After, the fiber was exposed to the headspace for 50 min at room temperature, the samples were desorbed in the injection port of GC-MS. To finalize the extraction, the SPME fiber is transferred to the injection port to separate the compositions of the samples.

### Gas chromatography-electron impact mass spectrometry (GC-EIMS) analysis

Fruit extract was phytochemical analyzed by GC-EIMS, using a Varian (Palo Alto, CA) CP 3800 gas chromatographs equipped with a capillary column DB-5 (30 m x 0.25 mm I.D., 0.25 µm film thickness; Agilent) to separate extract components. The interface for splitting the GC eluent to EIMS and MS detectors was custom designed and made by Agilent Technologies, and 2000 Varian Saturn trap mass ion detectors were used for the detection of fragment ions of derivatives from sesquiterpenes and non-terpenes. A commercial heated GC transfer line (Agilent Technologies) was used to connect the GC to MS. GC conditions were tested to find a good compromise between the analysis time and the resolution. The injection and transfer line temperatures were 250 and 240 ºC, respectively. The oven temperature was programmed as follows: 60-240 ºC at 3 ºC/min; helium was used as carrier gas (1 mL/min). The identification of the compounds is based on a comparison of the retention times with those of the authentic samples by comparing the linear retention indices (LRI) with respect to a series of n-hydrocarbons, as well as on the computer corresponding to the commercial (NIST 98 and Adams) and data from library mass spectra house literature, and MS [21]. In addition, the molecular weights of all identified substances were confirmed by gas chromatography with chemical Ionization mass spectrometry (GC-CIMS), using methanol as ionizing gas.

### Cell culture

The pancreatic β cell line INS-1E was cultured in a humidified chamber with 5% CO_2_ in RPMI-1640 medium (Merck, Sigma Aldrish, Darmstadt-Germany) at 11.1 mM glucose supplemented with 5% (vol./vol.), heat-inactivated foetal bovine serum, 10 mM HEPES, 2 mM L-glutamine, 100 U/mL penicillin, 100 μg/mL streptomycin, 1 mM sodium pyruvate and 50 μM β-mercaptoethanol. All experiments were performed in cells from passages 40 to 50. After 4 days of pre-culture in a T75 flask, cells were transferred to T25 flasks or 96-well plates and maintained 2 days at 11.1 mM glucose for further use.

### Cell viability assay

Crystal violet staining assay was used to analyzed cell viability of the PL extracts. Cells were seeded at low density (10^4^ cells/well) in 96-well plates for 24□h. Cells were treated with concentrations of 0.03215, 0.0625, 0.125, 0.25, 0.5 and 1 mg/mL of PLE dissolved in PBS, for 24□h and 72□h. At the end of the treatment, cells were washed with phosphate-buffered saline (PBS), stained for 5□min with a crystal violet solution [0.5% (w/v) crystal violet in 25% (v/v) methanol], and then gently rinsed with water. Absorbance was measured spectrophotometrically at 570□nm in a Specttra max molecular devices 384 plus plate reader. Data was expressed as the percentage of control.

### Treatments

Upon 85% cell confluency, cells in T25 flask were treated with glucose and/or plant extracts to determine the gene expression of insulin- and inflammatory-related genes. After 2 days of cell cultures in RPMI growth medium, cells were exposed to high glucose concentration (18 mM) in the presence or absence of the four extracts (0.125 mg/mL) during 24h. Then, cells were washed twice and pre-incubated for 30 min at 37 ºC in glucose-free Krebs-Ringer bicarbonate HEPES buffer (KRBH), as an insulin carrier [22]. The composition of the KRBH was 135 mM NaCl, 3.6 mM KCl, 5 mM NaHCO_3_, 0.5 mM NaH_2_PO_4_, 0.5 mM MgCl_2_, 1.5 mM CaCl_2_, and 10 mM HEPES, pH 7.4. BSA (0.1%). After one wash in glucose-free KRBH, cells were treated for 30 min in KRBH with high glucose concentration (18 mM) in the presence or absence of the four extracts (0.125 mg/mL). Treatments were stopped on ice. Cells were collected, frozen, and stored at -80°C until used.

### RNA isolation, reverse transcription and quantitative RT-PCR

INS-1E cells were lysed in TRIzol (Sigma-Aldrich, St. Louis, USA), and the total RNA fraction was extracted following the manufacturer’s instructions. Total mRNA was reverse-transcribed by the NZY First-Strand cDNA Synthesis Kit (NZYTech, Lda. Genes and Enzymes, Lisboa, Portugal). RT-qPCR was performed in a QuantStudioTM 5 Real-Time PCR System (Applied Biosystems, Massachusetts, USA). Genes were amplified employing the AceQ ® qPCR SYBR Green Master Mix (NeoBiotech, Nanterre, France) following the manufacturer’s instructions. The mRNA level of housekeeping gene cyclophilin A was used as an internal control for the normalization of the analyzed genes. All the RT-qPCR runs included non-template controls (NTCs). Experiments were performed in triplicates. Quantification of expression (fold change) from the Cq data was calculated by the QuanStudioTM Design & Analysis Software (Applied Biosystems). Details of the nucleotide sequences of the used primers are detailed in (Table 1).

**Table 1.**
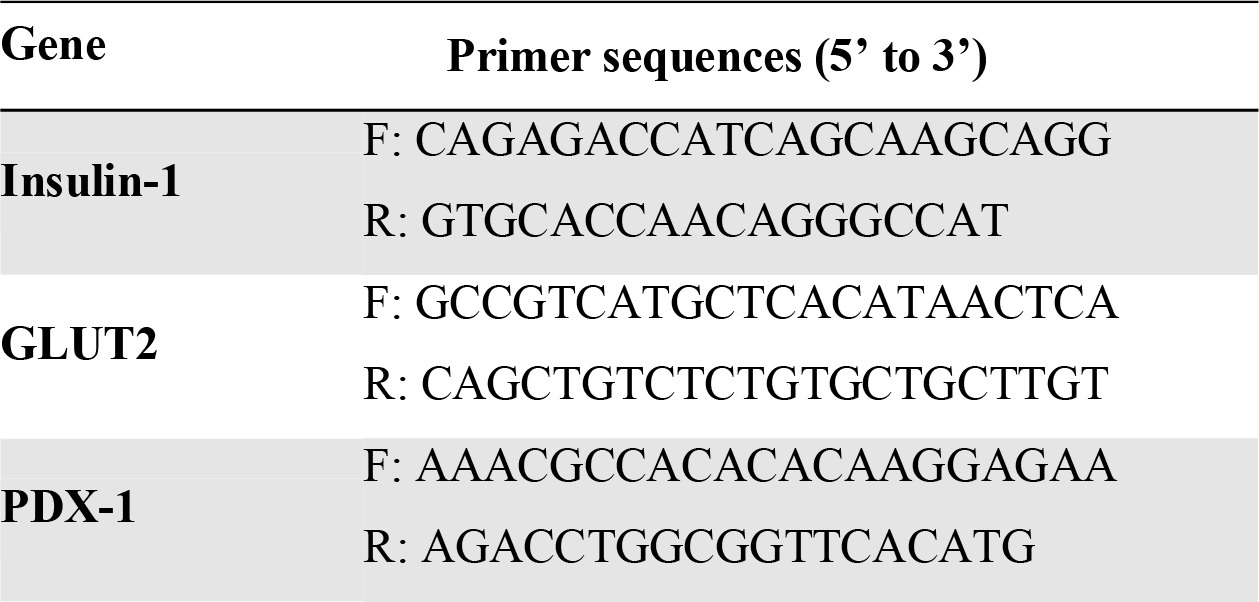
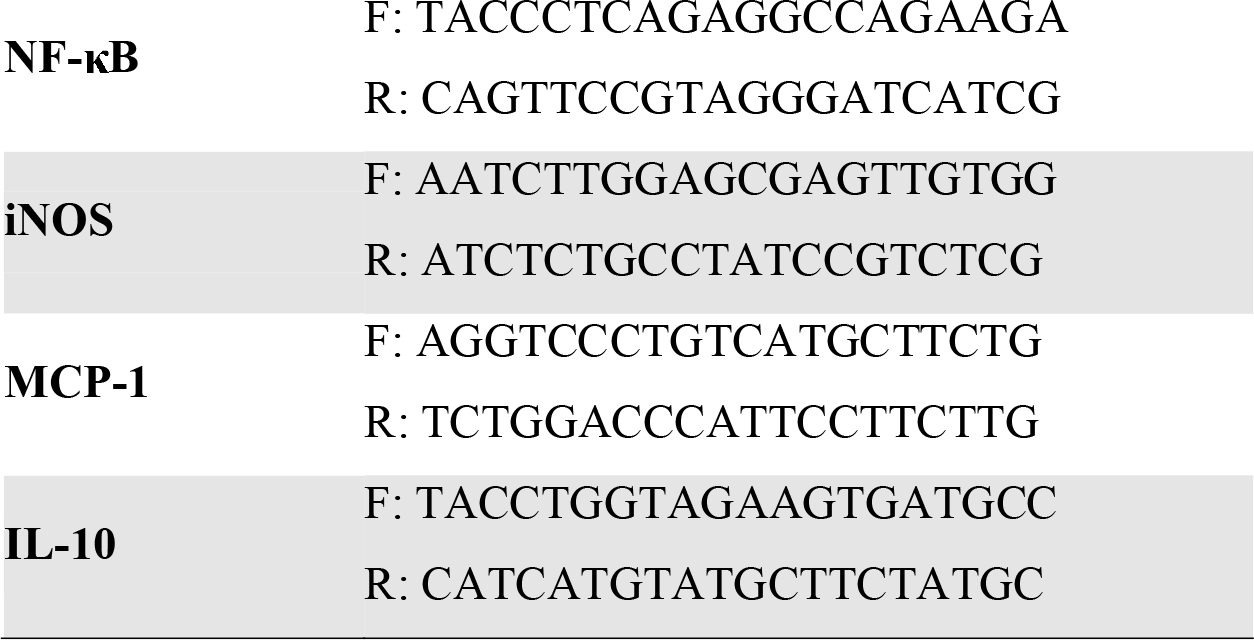
Nucleotide sequences of the primers used for the RT-PCR of genes.

### Statistical analysis

The results are reported as the mean±SEM. All the statistical parameters used were calculated with SPSS v28. The Shapiro-Wilk test was used to test for data distribution normality. To establish differences between experimental conditions, we used Student’s *t* test and one-way ANOVA followed by the Bonferroni post hoc test. Mann-Whitney U test and Kruskal-Wallis test were used as non-parametric alternatives, respectively. Values of p ≤ 0.05 were considered statistically significant.

## RESULTS

### Analysis of the chemical composition of *Periploca laevigata* extracts

The phytochemical screening revealed that PL contains tannins (hydrolysable and condensed), several types of flavonoids (flavones in root, flavanols in fruit and flavanones in leaves), and cardiac glycosides in root. However, alkaloids and anthraquinones were not detected in this study (Table 2).

**Table 2.** Major chemical groups identified in fruit extract of P. Laevigata.

| Chemical groups | Organ |  |  |
| --- | --- | --- | --- |
|  | Root | Fruit | Leave |
| Hydrolysable tannins | +++ | +++ | +++ |
| Condensed tannins | ++ | ++ | ++ |
| Flavones | ++ | - | - |
| Flavonols | - | +++ | - |
| Flavanones | - | - | ++ |
| Anthraquinones derivatives | - | - | - |
| Modified anthraquinones<br>(c.glycoside) | - | - | - |
| Cardiac glycosides | ++ | - | - |
| Alkaloids | - | - | - |

Footnotes: -, no coloring; ++, medium coloring; +++, intense coloring.

GC-EIMS and SPME-GC-MS analyses revealed the presence of 18 identified compounds. Table 3 showed that samples were rich in sesquiterpene hydrocarbons (61.9%), non-terpene derivatives (32.4%) and apocarotenes (5.1%). The most predominant components of sesquiterpene hydrocarbons were γ-muurolene (1478 LRI), α-copaene (1377 LRI) and δ-cadinene (1524 LRI) with a total amount of 29.1, 13.9% and 12.9%, respectively. Notably, it has been described anti-diabetic and anti-inflammatory functions for tannins, flavonoid and sesquiterpene hydrocarbons [23,24].

**Table 3.** Concentration of major secondary metabolites found in fruit extract of PL analyzed by SPME-GC-MS and GC-ECD techniques.

| Constituents | LRI | Powder fruit |
| --- | --- | --- |
| 2-pentyl furan | 993 | 4.5 |
| (Z)-3-octenal | 1036 | 6.1 |
| 4-methyldecane | 1059 | 1.7 |
| *n*-undecane | 1100 | 1.1 |
| Nonanal | 1102 | 3.5 |
| (E)-2-nonenal | 1163 | 3.4 |
| Naphthalene | 1181 | 4.9 |
| *n*-dodecane | 1200 | 1.8 |
| Decanal | 1205 | 1.6 |
| *n*-tetradecane | 1400 | 1.4 |
| *n*-pentadecane | 1500 | 2.4 |
| Non-terpene derivatives |  | 32.4% |
| α-copaene | 1377 | 13.9 |
| β-caryophyllene | 1419 | 3.9 |
| γ-muurolene | 1478 | 29.1 |
| Viridiflorene | 1495 | 1.1 |
| δ-cadinene | 1524 | 12.9 |
| α-calacorene | 1543 | 1.0 |
| Sesquiterpene hydrocarbons |  | 61.9% |
| (*E*)-geranylacetone | 1456 | 5.1 |
| Apocarotenes |  | 5.1% |
| Number of total compounds |  | 18 |

LRI: linear retention indices.

### *Periploca laevigata* extracts potentiate the expression of insulin-related genes in glucose-treated INS-1E cells

Considering the beneficial roles of medicinal plants to control the diabetic complications of T2D, this study evaluates the anti-diabetic functions of PL extracts in the pancreatic β cell line INS-1E. Firstly, we analyzed the cell viability of the PL extracts used from root, fruit and leaves (PLR, PLF, PLL and PLL-E) in INS-1E cells. After 24-hour incubation period, PLR and PLF extracts showed similar cell viability than control cells, whereas PLL and PLL-E extracts increased the cell viability with respect to control cells. Notably, cell incubated with the PLL-E concentration of 0.125 and 0.5 mg/mL revealed significant increases in the cell viability when compared with control cells (Fig. 1A). In addition, all PL extracts (PLR, PLF, PLL and PLL-E) after 72 hours of incubation induced an increase of the cell viability with respect to untreated cells (Fig. 1B). PLF extract at 0.0625 and 0.125 mg/mL and in PLL-E extract at 0.03215 and 0.0625 mg/mL exhibited significantly differences in the cell viability after 72 hours of incubation with respect to the control cells. Since no cytotoxicity was detected in cells treated for 24 hours with 0.125 mg/mL of PL extracts, this concentration was used to carry out the cell culture assays.

**Figure 1.**
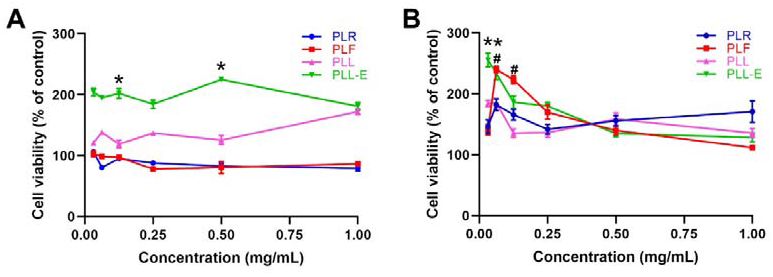
Cell viability analysis of INS-1E cells treated with increasing concentrations (0.03215, 0.0625, 0.125, 0.25, 0.5 and 1 mg/mL) of the different PL extracts for 24 **(A)** and 72 **(B)** hours. PL extracts were used from root (PLR), fruit (PLF) and leaves (PLL and PLL-E). Data was expressed as the percentage of control (mean ± SEM, n = 4 samples/group). * p < 0.05, PLL-E compared to the control group; # p < 0.05, PLF compared to the control group.

Then, we analyzed the effects of the PL extracts on insulin gene expression in glucose-treated INS-1E cells. Cells were stimulated with high levels of glucose concentration (18 mM) during 24 hours. Figure 2 demonstrated that the incubation of INS-1E cells with PLR, PLL and PLL-E extracts induced high upregulation of the insulin gene expression with respect to control cells, although these increases were not significant.

**Figure 2.**
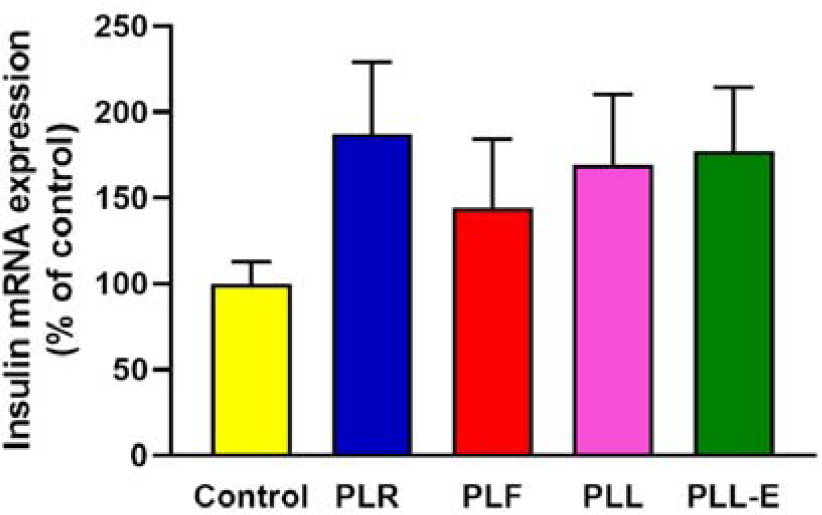
The PL extracts potentiates the expression of insulin gene in INS-1E cells treated with 18 mM of glucose during 24 hours. The mRNA levels were analyzed by qRT-PCR and cyclophilin A was used as the housekeeping gene. PL extracts were used from root (PLR), fruit (PLF) and leaves (PLL and PLL-E). Data represent mean ± SEM, n = 4 samples/group.

To gain better insight into the role of PL extracts in the insulin activity of glucose-treated INS-1E cells, the expression of the GLUT-2 glucose transporter was also evaluated. Figure 3 showed that the PLL-E extract increased significantly the GLUT-2 gene expression in glucose-treated INS-1E cells with respect to control cells. In addition, the PLR and PLF extracts exhibited a tendency to increase the GLUT-2 gene expression in the cells treated with glucose.

**Figure 3.**
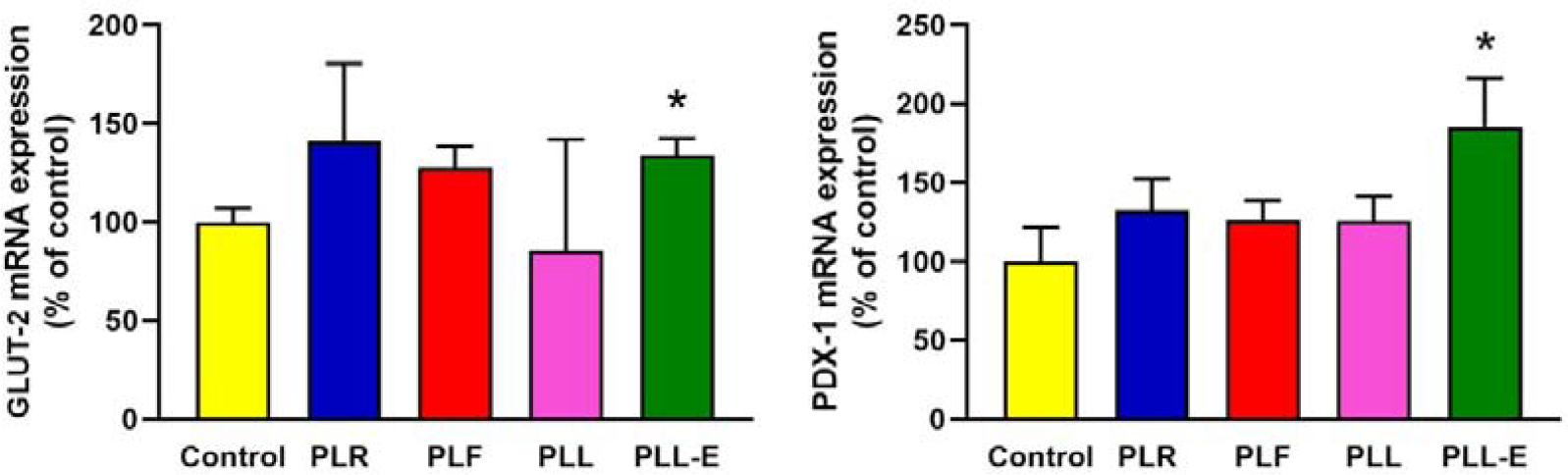
The PL extracts potentiates the expression of insulin-related genes in INS-1E cells treated with 18 mM of glucose during 24 hours. The mRNA levels of GLT-2 and PDX-1 were analyzed by qRT-PCR and cyclophilin A (Ppia) was used as the housekeeping gene. PL extracts were used from root (PLR), fruit (PLF) and leaves (PLL and PLL-E). Data represent mean ± SEM, n = 4 samples/group. * p < 0.05, compared to their respective control group.

Since that the transcription factor PDX-1 directly binds to the insulin gene promoter, regulating the mature β-cell function [25], the gene levels of this transcription factor were also analyzed in INS-1E cells treated with glucose and PL extracts. Our results revealed that PLL-E extract was capable to induce a significant gene expression of PDX-1 in glucose-treated INS-1E cells when compared with untreated cells (Fig. 3).

### *Periploca laevigata* extracts potentiate the expression of inflammatory-related genes in glucose-treated INS-1E cells

Several studies report that an inflammatory response occurs as a result of the activation of the immune response to high blood glucose levels [26]. Considering that PL extracts show anti-inflammatory and anti-oxidant functions [18], we also evaluated the role of the four PL extracts (PLR, PLF, PLL and PLL-E) in the expression of inflammatory genes (NF-κB, iNOS, MCP-1 and IL-10) in glucose-treated INS-1E β cells. Figure 4 showed that the PLF and PLL extracts decreased significantly the iNOS gene expression in glucose-treated INS-1E cells with respect to control cells. Although no significant differences were showed in the NF-κB and MCP-1 gene expressions, all PL extracts exhibited a marked tendency to decrease both gene expressions in glucose-treated cells. Notably, the PLR, PLF and PLL-E extracts increased significantly the gene expression of the anti-inflammatory cytokine IL-10 in glucose-treated INS-1E cells with respect to control cells (Fig. 4).

**Figure 4.**
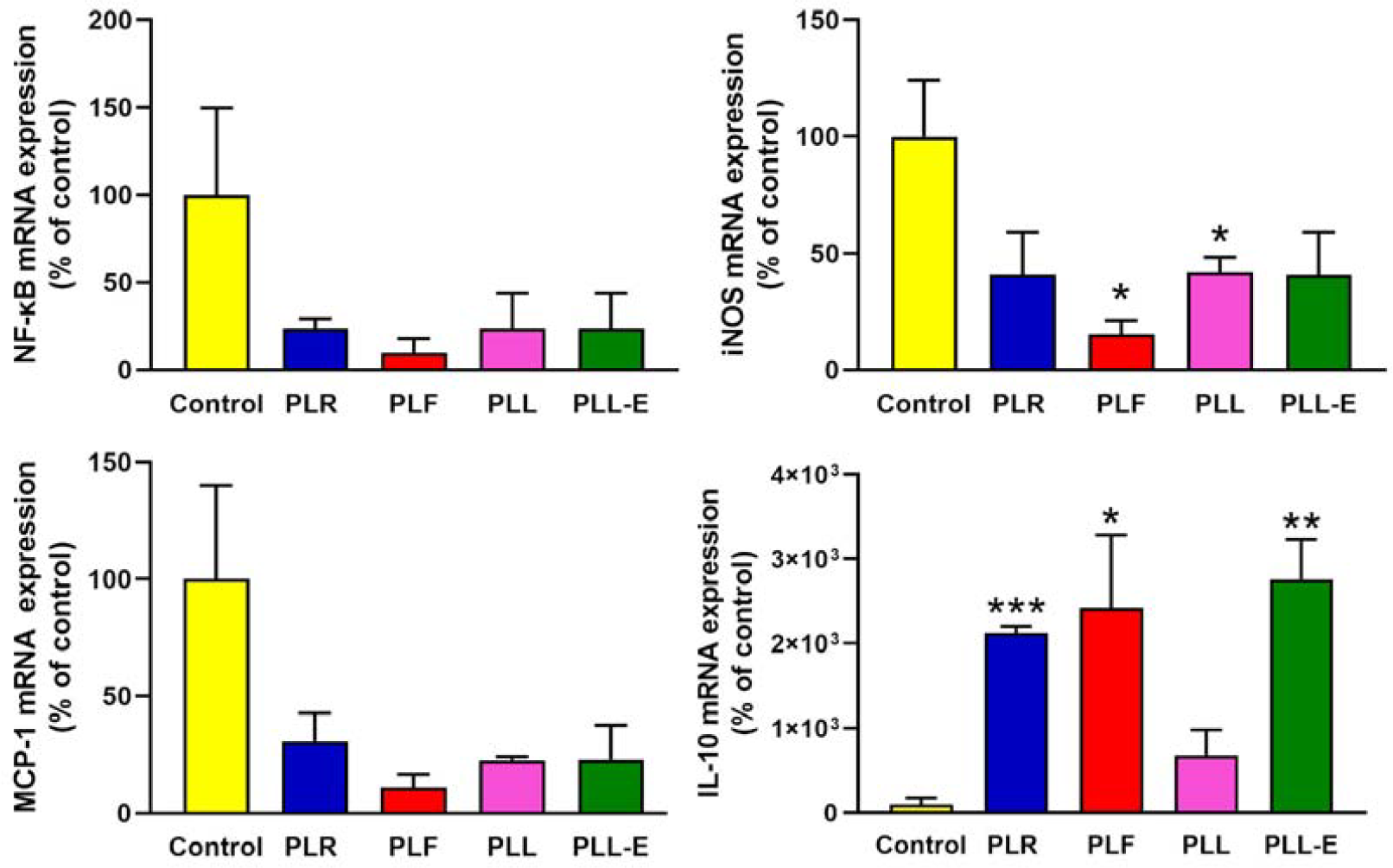
The PL extracts decrease the levels of inflammatory genes in INS-1E cells treated with 18 mM of glucose during 24 hours. The mRNA levels of NF-κB, iNOS, MCP-1 and IL-10 were analyzed by qRT-PCR and cyclophilin A was used as the housekeeping gene. PL extracts were used from root (PLR), fruit (PLF) and leaves (PLL and PLL-E). Data represent mean ± SEM, n = 4 samples/group. * p < 0.05, compared to their respective control group.

## DISCUSSION

It has been previously demonstrated the beneficial roles of medicinal plants to control the diabetic complications of T2D [14], and the therapeutic role of PL extracts [18]. In the present study, we report the potentially beneficial effects of some PL extracts on the expression of insulin- and inflammatory-related genes under high glucose conditions in the pancreatic β cell line INS-1E (Fig. 5). Overall, these findings suggest that PL may represent a useful therapeutic approach to treat the pancreatic β cell dysfunction and the inflammation associated with the T2D.

**Figure 5.**
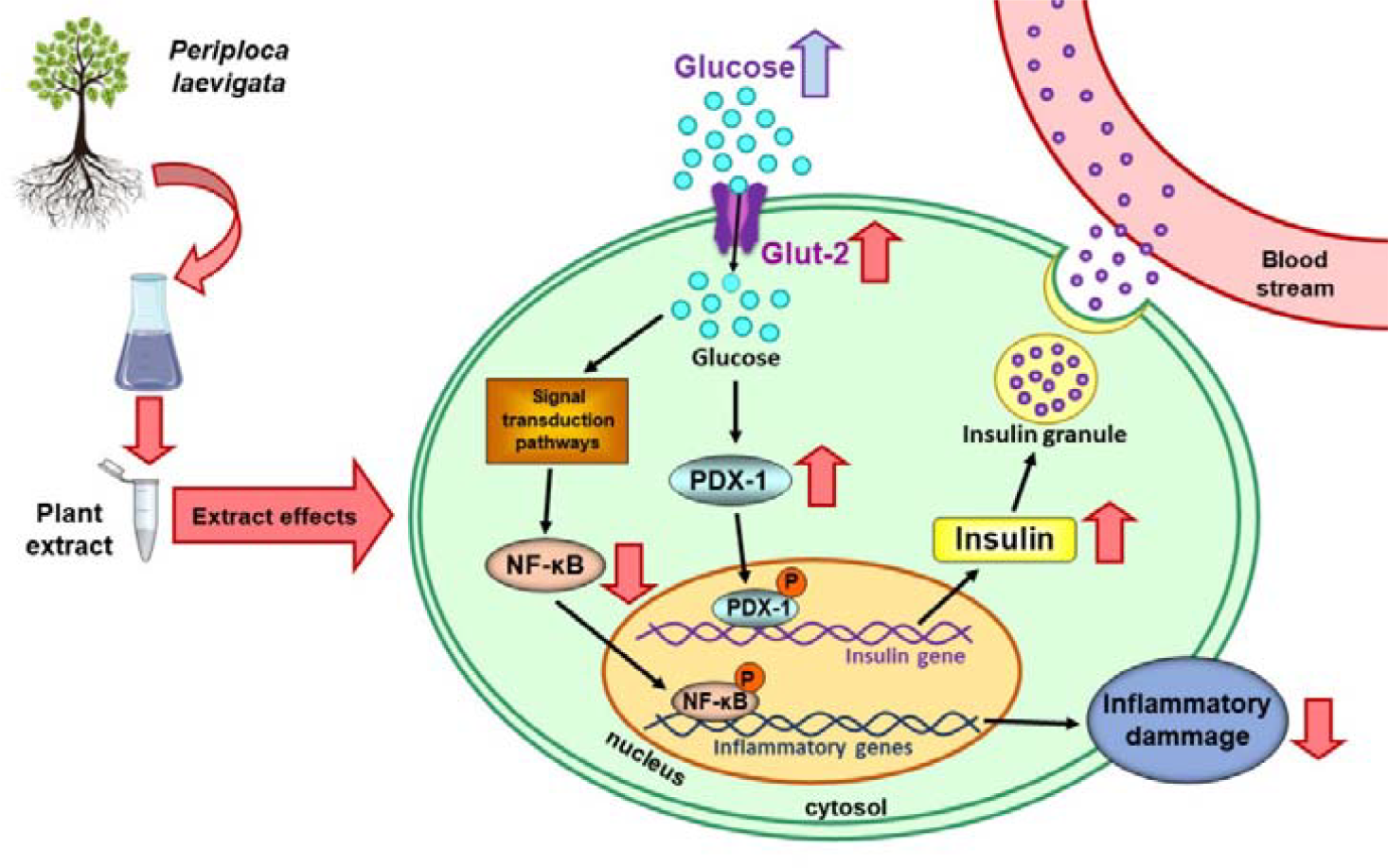
Schematic representation of the possible anti-diabetic and anti-inflammatory functions of *Periploca laevigata* extracts in pancreatic β cells in high blood glucose levels. PL extracts are able to increase the gene expression of insulin, GLUT-2 and the transcription factor PDX-1 in the presence of high glucose levels. In addition, PL extracts are also capable to decrease the gene expression of NF-κB, iNOS and MCP-1, as well as increase the anti-inflammatory cytokine IL-10 in the presence of high glucose levels.

The T2D is associated with pancreatic β cell dysfunction, which leads to insulin resistance and loss of insulin secretion [2]. It has been reported that the proinsulin synthesis and the β cell sensitivity to glucose, in terms of hormone secretion, appear to be modulated by insulin signaling. Indeed, alterations of the insulin signal transduction pathway could induce β cell dysfunction, contributing to the pathogenesis of T2D [27]. For instance, the pancreatic transcription factor PDX-1, by regulating the insulin promoter, plays a key role in pancreatic β cell function [28]. It has been shown that both glucose and insulin stimulate PDX-1 binding activity, and reductions in PDX-1 expression may impair glucose-stimulated insulin secretion [29]. Our results revealed that some PL extracts could increase the insulin and PDX-1 gene expression in a high-glucose condition in INS-1E β cells, suggesting a protective role of PL extracts in the diabetic complications. In this sense, previous studies have reported that a sylvestre plant extract was capable to stimulate insulin secretion at high glucose concentration on human islets both *in vivo* and *in vitro* [30]. Moreover, a polysaccharide from mulberry leaves on diabetic mice may modulate the hepatic glucose metabolism and gluconeogenesis, by up-regulating the PDX-1 and insulin expressions in pancreas [31].

The glucose uptake by pancreatic β cells is mediated by GLUT-2 receptor and its deficiency and reduction of its expression could cause T2D [32]. In our case, the transport of glucose is predominantly conducted by GLUT-2 in INS-1E cells, it allows a parallel rise of pancreatic β cell function in response to an increase of blood glucose concentration [33]. Indeed, it has been demonstrated that PDX1-GLUT-2 pathway may be one of the signaling pathways underlying pancreatic beta cell insulin resistance. Thus, the reduction of PDX-1 expression results in a decline in GLUT-2 expression [32]. Interestingly, our results demonstrated that PLL-E extract is able to upregulate the expression of PDX-1 and GLUT-2 genes at high glucose levels, in order to improve the pancreatic β function. Accordingly, hypoglycemic plant extracts are able to improve the expression of PDX-1 and GLUT-2 [33].

Recent studies reported that the inflammation due to the glucotoxicity contributes to the loss of metabolic tolerance and increases the risk of insulin resistance [34,35]. For instance, the increase expression of MCP-1 can induce insulin resistance and infiltration of macrophages into adipose tissue [36], and iNOS has been associated with the pathogenesis and progression of several diseases, including insulin resistance [37]. In fact, our results show that treatment of β cells with some PL extracts decreases significantly the gene expression of iNOS and there is a tendency to decrease in MCP-1 and NF-κB gene expressions. In relation to this, it has been shown that inhibition of NF-κB activation can protect pancreatic β cells from cytokine-induced apoptosis. This inhibition could potentially be considered as an effective approach to protecting β cells and the prevention of diabetes by regulation of production of multiple proinflammatory cytokines (TNF-α, IL-1β, IL-6 and IFN-γ), the chemokine MCP-1, and enzymes (iNOS and COX-2) [38]. On the other hand, our data also showed that treatment with PL extracts significantly increased the levels of the anti-inflammatory cytokine, IL-10, at high glucose conditions, suggesting a possible protective role of PL against the inflammatory state in T2D. Similarly, IL□10 gene transfer reduced the expression of inflammatory cytokines, attenuated pancreatic insulitis and inhibited β cell apoptosis [39]. In addition, anti-inflammatory molecules, such as IL-10, IL-4 and IL-13, can exert a direct influence in the β-cell function and viability and the circulating levels of these cytokines may be reduced in diabetes [40].

It has been reported that the therapeutic actions of plant extracts are associated to their composition in some secondary metabolites. In our case, the anti-diabetic and anti-inflammatory functions of PL extracts in glucose-treated pancreatic β cells may be due to the high abundance of tannins and flavonoids, which have shown anti-diabetic, anti-oxidative, cardioprotective, neuroprotective, anti-inflammatory effects [23,24,41]. Furthermore, sesquiterpene hydrocarbons also showed anti-diabetic and anti-inflammatory functions [23,24]. Natural constituents from PL such as proteins, phenolic compounds and polysaccharide have a preventive potential and *in vivo* treatment of inflammation [17]. According to the same plant family, *Apocynaceae*, also showed anti-diabetic effect [42]. Regarding the polyphenols, which are present in our extract in high abundance, are able to improve the gene expression of PDX-1 and GLUT-2, to ameliorate pancreatic function [43].

In summary, our results suggest that PL can be exerted as anti-diabetic and anti-inflammatory functions against the T2D, without inducing pancreatic β cell toxicity. These findings provide evidence of the therapeutic potential of PL extracts and may provide a new means to treat the diabetic dysfunction and pancreatic inflammation associated with the diabetes.

## ACKNOWLEDGMENTS

This work has been supported by grants from the Spanish Ministry of Health[PNSD (2019□I039), GVA (CIAICO/2021/203), the Primary Addiction Care Research Network (RD21/0009/0005) and FEDER Funds, GVA and the Tunisian ministry of Higher Education and Scientific Research.

## AUTHOR CONTRIBUTIONS

Conceptualization, G.T., LCG. and M.P.; methodology, G.T., S.M. and M.P.; formal analysis, G.T. and S.M.; investigation, G.T., S.M. LCG and M.P.; resources, M.P.; data curation, S.M. LCG. and M.P.; writing-original draft preparation, G.T., S.M. and M.P.; writing-review and editing, G.T., S.M. and M.P.; supervision, L.C.G. and M.P.; funding acquisition, M.P, LCG. All authors have read and agreed to the published version of the manuscript.

## DATA AVAILABILITY STATEMENT

The data that support the findings of this study are available on request from the corresponding author. The data are not publicly available due to privacy or ethical restrictions.

## ABBREVIATIONS

GLUT-2: Glucose transporter 2
iNOS: Inducible nitric oxide synthase
MCP-1: Monocyte chemoattractant protein-1
NF-kB: Nuclear factor kappa B
PDX-1: Pancreatic duodenal homeobox 1
PLF: Periploca laevigata fruit
PLL: Periploca laevigata leaves
PLL-E: Periploca laevigata leaves in hydro-ethanolic solution
PLR: Periploca laevigata root

